# Assessment of resting state structural-functional relationships in perisylvian region during the early weeks after birth

**DOI:** 10.1101/2023.10.26.564007

**Authors:** Roxana Namiranian, Hamid Abrishami Moghaddam, Ali Khadem, Reza Jafari, Aimirhossein Chalechale

**Author notes:** **Corresponding Author:** Hamid Abrishami Moghaddam, Ph.D., Faculty of Electrical Engineering, K.N. Toosi University of Technology, PO Box 16315-1355, Tehran, Iran. **E-mail addresses:** Roxana Namiranian, Ali Khadem, Reza Jafari, Amirhossein Chalechale.

## Abstract

This study investigates the structural-functional (S-F) relationships of the perisylvian region during the first weeks after birth in the resting state. Previous joint S-F studies of perisylvian development were mainly conducted on individual structural and functional metrics. By employing a weighted combination of metrics, joint S-F studies can enhance the understanding of perisylvian development in neonates, thereby providing valuable features for the prediction of neurodevelopmental disorders. To this end, we employed both structural and functional metrics derived from 16 perisylvian sub-regions (PSRs). Structural metrics included morphological and myelination measures, while functional metrics encompassed functional connectivity (FC) values between a designated seed PSR and the remaining PSRs. In addition, fractional amplitude of low-frequency fluctuations (fALFF) for all PSRs was computed and used as a distinct group of functional metrics. During statistical procedure, based on sparse canonical correlation analysis (CCA), the structural metrics were correlated with each respective group of functional metrics. Then, CCA was employed to delineate regional interdependencies and derive combined structural-functional features. The findings revealed that combined features outperform individual metrics for characterizing the normal development of the PSRs in term neonates. The myelination is prominently related to both fALFF and FCs, while morphological metrics of the PSRs have very limited contribution in combined structural features. Among the designated PSRs, the FCs of both the right insula and Heschl’s gyrus with other PSRs demonstrated a more robust correlation with the combined structural features.

## 1. Introduction

The perisylvian region of the brain plays a crucial role in processing the language and auditory communication, making it an area of significant interest for researchers studying early brain development. Examining the structural-functional (S-F) relationships in perisylvian sub-regions during the early postnatal period offers critical insights into the underlying mechanisms of infant language and auditory development. The development of advanced neuroimaging methodologies, such as magnetic resonance imaging (MRI) and functional MRI (fMRI), has provided researchers with non-invasive tools to assess the growth of neonatal brain structure and function. Through integrating data from both structural and functional neuroimaging, researchers can achieve a more comprehensive understanding of how the anatomical organization of the brain relates to its function during this critical developmental phase.

The postnatal period coincides with extremely dynamic and rapid brain changes [1,2], which are crucial for understanding how brain evolution influences lifelong cognitive abilities and behaviors [3,4]. This period is also important for early diagnosis or even prognosis of neurodevelopmental disorders [5–7]. However, in this period, multimodal (e.g., structural-functional) brain data acquisition and analysis are challenging [8,9] due to neonates’ non-cooperativeness, immature brain structure, and inability to respond deliberately to task-based stimuli. Nevertheless, recent advances in neuroimaging and open resource projects such as the developing Human Connectome Project (dHCP) [10] have provided researchers with suitable multimodal datasets for structural and functional developmental studies [11–17].

Language and auditory related sub-regions were also of particular interest in most of the postnatal neurodevelopmental studies, since they can be performed without requiring a deliberate response of the subjects [18]. Accordingly, task based or resting-state functional studies have been performed to examine language and auditory sub-regions and their interconnections with other functional circuits [19–22]. The relationship between functional connectivity (FC) and both language [23,24] and auditory [25–28] performance has been demonstrated using resting-state fMRI (rs-fMRI). Xiang et al. [29] showed that the topology of FC in six frontal sub-regions of the perisylvian language network during rest is consistent with the topology of language network of the brain. Nevertheless, few researchers have investigated the association between the structure and function of auditory- and language-related regions during rest in early childhood [30–32]. Furthermore, when comparing resting-state studies to task-based studies, the data acquisition procedure is typically easier, resulting in larger datasets. Consequently, the outcomes and models derived from resting-state studies are deemed more reliable.

A few task-based studies have attempted to examine structural features to correlate with functional performance in language and auditory regions. Daneshvarfard et al. [33] correlated a microstructural metric of the superior temporal gyrus (STG) with latency of auditory response in 17 preterm neonates with gestational age (GA) between 28.4 and 32.2 weeks. Moreover, Adibpour et al. [34] demonstrated that the microstructural measures of the inferior frontal gyrus (IFG) modulate the auditory responses in the first 6 months after birth, even after controlling for age. Studies conducted in the first postnatal semester demonstrated a correlation between the myelination of corpus callosum fibers and speed of inter-hemispheric auditory and visual information transfer [34–36]. These studies assert the importance of cortical properties in identifying specialized structural features associated with auditory function [34]. Morphological metrics like surface area and folding measures, before term equivalent age, are related to the later language outcome [4,37]. Furthermore, discrepancies are evident in recent whole-brain resting-state studies. In term neonates, cortical thickness has been reported to correlate with the anterior-to-posterior gradient of resting-state FCs [11]. However, for both term and preterm individuals, after controlling for age, no significant associations were observed between morphological characteristics, local functional activity, or resting-state FCs [16]. To our knowledge, no study has been conducted to determine whether shape-related cortical metrics are significantly correlated to functions of auditory or language-related regions, during the first few weeks after birth.

Previous neonatal joint S-F studies on language and auditory regions investigated the relationship between individual functional and structural metrics [33,34,36]. They did not explore whether a combination of structural metrics might be associated with a mixture of functional measures. For example, the joint study in [33], showed that maturation of the auditory responses is correlated with myelination of auditory pathways and regions, while they become uncorrelated after controlling for age. Another joint analysis [34] failed to find the link between functional lateralization in auditory responses and structural asymmetries, while asymmetric maturation of perisylvian sub-regions (PSRs) and pathways [38,39] was consistent with the language and auditory laterality [40–43]. These findings are in accordance with the fact that there are complex relationships between brain structure and function [44], and that more informative structural and functional features are required to comprehend the links. Combined features extracted from structural and functional domains respectively, are hypothetically more promising than individual metrics for understanding complex S-F couplings.

It is worth noting that joint studies seek to identify comprehensive structural features that can better predict brain function [11,17,34]. Such features can be used effectively for early diagnosis at ages when symptoms of cognitive impairments have not yet appeared or functional measures are difficult to acquire. While joint S-F analysis facilitated the diagnosis of language and neurodevelopmental disorders during adulthood [45–47], such joint studies have not yet been conducted on neonates. Zhang et al., [15] discovered atypical patterns of S-F couplings in premature birth with respect to term neonates. Further studies with a thorough investigation into combined and innovative structural and functional features, can provide comprehensive information [5,48] about the normal maturation of the brain.

To address the aforementioned issues, we utilized structural MRI and rs-fMRI datasets and adopted a feature extraction approach to investigate functional and structural features of language-related regions involved in the normal S-F relationships during the first few weeks after birth. Specifically, we focused on examining the fractional amplitude of low-frequency fluctuations (fALFF), FC patterns, and structural metrics of these regions during rest. The importance of using the structural MRI modality lies in the fact that this modality is more practical for neonates and clinical applications due to its lower cost and faster data acquisition. We also used canonical correlation analysis (CCA) [49], a data-driven multivariate method, to simultaneously capture comprehensive linear interactions between multiple structural and functional measures of the brain. Initially, the incorporation of sparsity terms in the sparse CCA algorithm enables effective handling of high-dimensional data, addressing concerns of overfitting and lack of interpretability that conventional canonical correlation analysis may suffer from. Consequently, features are extracted based on the association between the two modalities, promoting joint information extraction rather than treating each modality separately [11,15].

The paper is structured as follows: Section 2 presents the materials and methods. Section 3 showcases the results. Section 4 discusses the alignment of the achievements with previous relevant studies and limitations of this work. Finally, the concluding remarks are given in Section 5.

## 2. Materials and methods

### 2.1. Neonatal dataset

The neonatal neuroimaging dataset released by dHCP [50] was employed in this study. This dataset provides multi-modal neuroimaging (functional, structural and diffusion MRI) sessions comprising around 500 neonates aged from 24 to 45 weeks GA in the second release (see http://www.developingconnectome.org for more details). In our study, only term neonates with the birth age greater than 37 weeks GA were participated. Moreover, the subjects with sever head motion during fMRI scans [51], or missing one of the structural or functional data were excluded. Therefore, a final set of 166 sessions was enrolled to investigate the structural and functional relationships as illustrated in Fig. S1 in Supplementary Materials (81 girls and 85 boys, age-range: 38-44.7 weeks GA, mean age ± STD: 41.3±1.6 weeks GA). All neonates were scanned during natural sleep on a 3T Phillips Achieva with a dedicated neonatal imaging system and a neonatal 32 channel phased array head coil at the Evelina Neonatal Imaging Centre, St Thomas’ Hospital, London. Hearing protection was provided for infants.

Anatomical MRI was acquired using a turbo spin echo (TSE) sequence for T2w and an IR (inversion recovery) TSE sequence for T1w images, in sagittal and axial slice stacks (in-plane resolution 0.8 × 0.8 mm^2^ and 1.6 mm slices overlapped by 0.8 mm). Other parameters were– T2w: TR/TE = 12000/156 ms; T1w: TR/TI/TE = 4795/1740/8.7 ms. Final resolution of reconstructed structural images, using slice-to-volume registration was 0.5L×L0.5L×L0.5 mm^3^ (for more details check [52]).

Resting-state fMRI using multiband (BM) 9× accelerated echo-planar sequence with 2.15 mm isotropic resolution (TE/TR = 38/392 ms) was performed for infant brain during 15 min. In addition to the main EPI scans, single-band EPI reference scans were also obtained with a readout bandwidth matching the main scans. Additional spin-echo EPI scans were acquired with phase-encoding directions of 4xAP and 4xPA. The reconstructions were performed using the extended SENSE framework described by Zhu et al. [53], with sensitivity maps calculated from the corresponding single-band data. Field maps were obtained using an interleaved spoiled gradient-echo sequence with dual TE (TR= 10 ms, TE1= 4.6 ms, TE2=6.9 ms, a flip angle =10°), an in-plane resolution of 3 mm isotropic, and a slice thickness of 6 mm (see [54] for more details).

### 2.2. The proposed pipeline

The proposed pipeline for extracting and assessing the interplay between structure and function of the neonatal brain is shown in Fig. 1. First, in the data processing and feature generation block, the brain is parcellated into 90 regions using the UNC (also known as neonatal AAL) atlas [55] from which 16 PSRs are selected for further processing. For all PSRs, four morphological attributes and one myelination index were derived and employed as 80 structural metrics. Subsequently, the FCs between a designated PSR and remaining PSRs were computed as functional metrics, which were then correlated with the structural measures. Additionally, the fALFF within all PSRs was calculated and employed as 16 functional measures to be correlated distinctly with the aforementioned structural metrics. Structural and functional metrics are employed to extract combined structural and combined functional (CS-CF) features. Initially, because of the collineality and high-dimensional dataset, we utilize the sparse CCA technique to select relevant features. Subsequently, we apply CCA to identify the most correlated combination of the selected structural and functional features. To evaluate CS-CF features as well as the canonical weights, the statistical analysis was employed on both canonical techniques. Finally, features with significant and stable statistical results were extracted from the CCA technique. The subsequent sections will delve into the details of this pipeline.

**Fig. 1.**
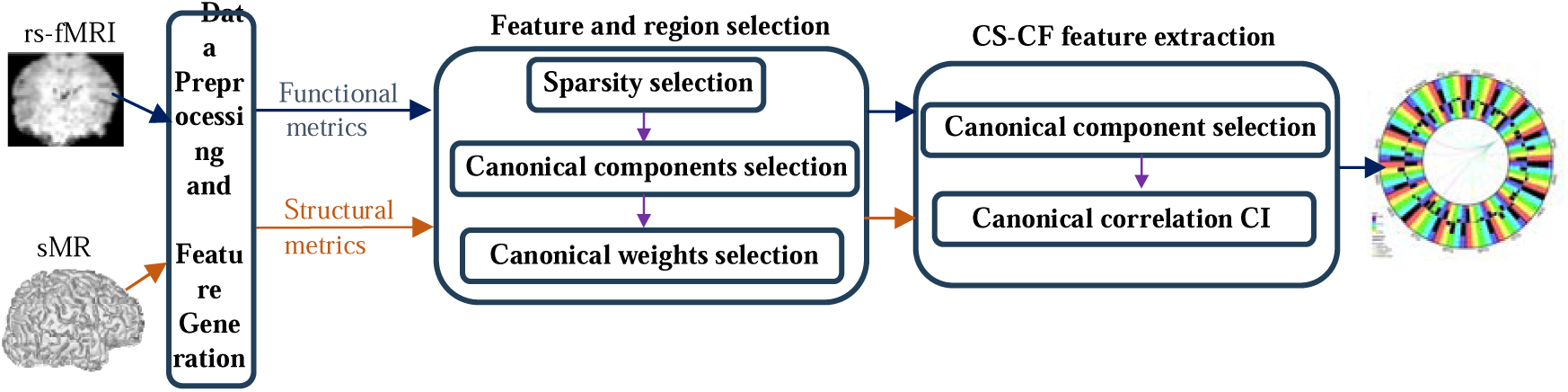
The proposed pipeline for extracting combined structural and functional features. Initially, in the data preprocessing and feature generation stage, structural and functional metrics are generated from structural MRI and rs-fMRI data, respectively. Structural metrics including morphological and myelination measures are generated for 16 PSRs which are designated on the UNC atlas. Two groups of functional metrics including the fALFF within PSRs and FCs between a designated PSR and remaining PSRs are generated and used separately for analysis. In the second stage, the sparse CCA technique is employed to select features and regions based on significant canonical components. Non-zero canonical weights are identified using a bootstrapping test. Next, the CS-CF features are extracted through canonical correlation CI of the first significant component of standard CCA. Finally, the combined structural and functional features (FCs) are visualized using a connectogram. sMRI: structural MRI, CS: combined structural, CF: combined functional, CI: confidence interval

#### 2.2.1. Dataset processing and feature generation

The neonatal brains were parcellated to 90 regions using UNC atlas [55]. It is a volumetric infant brain atlas proposed for 3 age-ranges including neonates, 1- and 2-years old infants. This atlas provides finer parcellation around Sylvian fissure compared to the atlas employed by dHCP pipeline. Among 90 parcels of UNC, the 16 PSRs are as follows: triangular gyrus (trianG), opercular gyrus (operG), supramarginal gyrus (suprG), angular gyrus (AG), STG, medial temporal gyrus (MTG), Heschl’s gyrus (HG), and insula in both hemispheres. The labelled parcels of UNC atlas were propagated to the 40-week templates based on multi-channel registration [52]. Age-specific UNC parcels were linearly and then nonlinearly registered to the structural space defined by T2w image of each participant. Volumetric parcel masks were used for functional data processing. The registered mask of each UNC parcel was mapped to the neonate’s cortical surfaces using Connectome Workbench from HCP tools (https://www.humanconnectome.org/software/connectome-workbench), and served for structural processing.

##### Structural data processing

In our study, we utilize four surface metric maps provided for each neonate by the dHCP pipeline [52]. The pipeline can be summarized as follows: the structural MRIs undergo some preprocessing steps, including motion and bias correction, and brain extraction. Subsequently, the cortical surfaces of each neonate are extracted and employed to estimate one microstructural and three morphological metrics. As a microstructural metric, the myelination index is estimated by the T1w/T2w ratio. As morphological metrics, the mean curvature, sulcal depth, and cortical thickness are computed. The surface metric maps provided by the dHCP team are averaged over the vertices of each parcel in surface registered UNC atlas. The surface area of each PSR is another structural measure that was computed to model the inter-subject variations of neonatal brains. As a result, structural metrics consist of 80 measures (5 structural measures of 16 PSRs) from 166 neonates.

##### Functional data processing

For the rs-fMRI, the preprocessing involved correction for geometric distortions and head motion, registration to T2w structural images, high-pass filtering (150 s), and ICA denoising [54]. The volumetric UNC labeled masks from native T2-space were transformed into native functional space (using transformation matrices provided by dHCP team). Then, FCs were calculated between each pair PSR using Pearson correlation [56,57] between regional time series extracted from 16 PSRs by averaging the blood-oxygenation-level dependent (BOLD) signal within each region. The fALFF of each PSR is calculated by integration of the power spectrum within the frequency range between 0.01 to 0.08 Hz relative to the integration of the power spectrum of full frequency range. We regressed out age from structural and functional metric sets to ensure that age, as a potential confounder of measures, does not drive the correlation.

#### 2.2.2. Feature and region selection

The interrelationships between structural and functional data were formally examined through the application of sparse CCA, with the objective of identifying the most relevant features and regions delineating this association. Sparse CCA, is a CCA algorithm adapted for high-dimensional data which incorporates feature selection by embedding lasso penalty in the CCA approach. Like CCA, which provides shared latent space (canonical variates) that maximizes the correlation between the two datasets, sparse CCA correlates datasets using sparse linear projection of both sets (canonical weights). Sparse CCA can be expressed as an optimization problem:

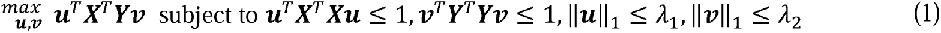

which finds canonical weights, ***u****_p_*_×1_ and ***v****_q_*_×1_ that maximize *cor*(***Xu***, ***Yv***). ***X****_n_*_×*p*_ and ***Y****_n_*_×*q*_ represent FCs of each PSR (*p* = 15) and structural metrics of UNC parcels (*q* = 80), respectively, with the same number of subjects in rows (n = 166). When fALFF is used as the functional metric, the corresponding dimension is *p* = 16. The sparsity is achieved by applying lasso penalties, which needs to be tuned using grid search between ranges of *λ*_l_ and *λ*_2_ parameters, from 0.1 to 0.98 in increments of 0.1. We employed the penalized multivariate analysis (PMA) package in the R programming language to implement the sparse CCA algorithm proposed by Witten et al. [58].

The determination of appropriate sparsity levels, denoted as *λ*_l_ and *λ*_2_ in sparse CCA formulation (1), has been a challenging task, extensively explored in previous studies[58–60]. They commonly performed permutation test for estimating the sparsity levels of canonical weights. In this approach, significance of each sparsity level in grid search is determined for the first canonical component, by a permutation test (*p* < 0.05) with 500 repetitions.

Subsequently, another permutation test (*p* < 0.05), also with 500 repetitions, is conducted to identify all other significant components at the selected sparsity level, based on the canonical correlation coefficient (CCC) of each component. Apart from selecting significant canonical components, examining the stable coefficients are also of great importance. Therefore, the next stage involves examining the coefficients using the bootstrap method in the sparse CCA, with 1000 resampling iterations. The obtained coefficients are subject to a statistical test to assess their stability. Coefficients with a 95% confidence interval (CI) overlapping with zero are considered as zero. Furthermore, the coefficient of variation is computed, and non-zero coefficients with a variation-to-mean ratio of less than 20% are regarded as appropriate.

#### 2.2.3. CS-CF feature extraction

We analyzed the selected features and regions derived from the stable coefficients of the significant components using sparse CCA. These features and regions were subjected to CCA, followed by a permutation test with 500 repetitions to validate the significant component. Benjamini-Hochberg false-discovery rate method was employed to correct for multiple comparison. Subsequently, for the first significant component, the bootstrping method with 1000 resampling iterations was employed to identify components that indicate acceptable S-F relations. Ultimately, the components with 90% CI of canonical correlation exceeding 0.3 were selected.

## 3. Results

This section presents the findings from two distinct experimental investigations. The first experiment investigated the relationship between the same 80 structural measures and 16 fALFF metrics derived from 16 PSRs (resulting in one metric per region). The second experiment examined the correlation between 80 structural measures obtained from 16 PSRs (entailing five metrics per region) and 15 FC metrics. These FC metrics were calculated between a designated seed PSR and the remaining PSRs. The objective was to identify combined structural and functional features that demonstrate the strongest relationship between structure and function.

In the second experiment, no significant sparsity level was found when operG, AG, MTG in both hemispheres were used as seeds for FC. Consequently, these regions were excluded from subsequent analysis in the second experiment. In both experiments, after bootstrapping and excluding metrics with zero values in their 95% CI, a limited number of metrics and PSRs survived for each significant component of sparse CCA. The results demonstrated that the structural measure strongly correlated with CF features is myelination. The sum of the absolute canonical weights for each metric, and the selected metrics and ROIs, are provided in Table S1 and Table S2 of Supplementary Materials, respectively. The selected metrics and ROIs were passed to CS-CF feature extraction step for further processing using CCA. The first significant component of CCA with CCC exceeding 0.3 based on 90% CI (Table S3 of Supplementary Materials) is selected as admissible CS-CF features. The first experiment which employed fALFF features resulted in an admissible CS-CF feature. In the second experiment, only when inula-R, HG-R and HG-L were used as seed regions for FC, their respective CS-CF features were admissible.

### 3.1. Structure-function relationship via individual versus combined features

The histograms of significant correlation coefficients between the individual structural and functional features corresponding to aforementioned admissible CS-CF features are shown in Fig. 2. The vertical red dotted line in each diagram specifies the correlation between extracted CS-CF features. As illustrated in Fig. 2, the combined features outperformed the individual ones in terms of correlation coefficient.

**Fig. 2.**
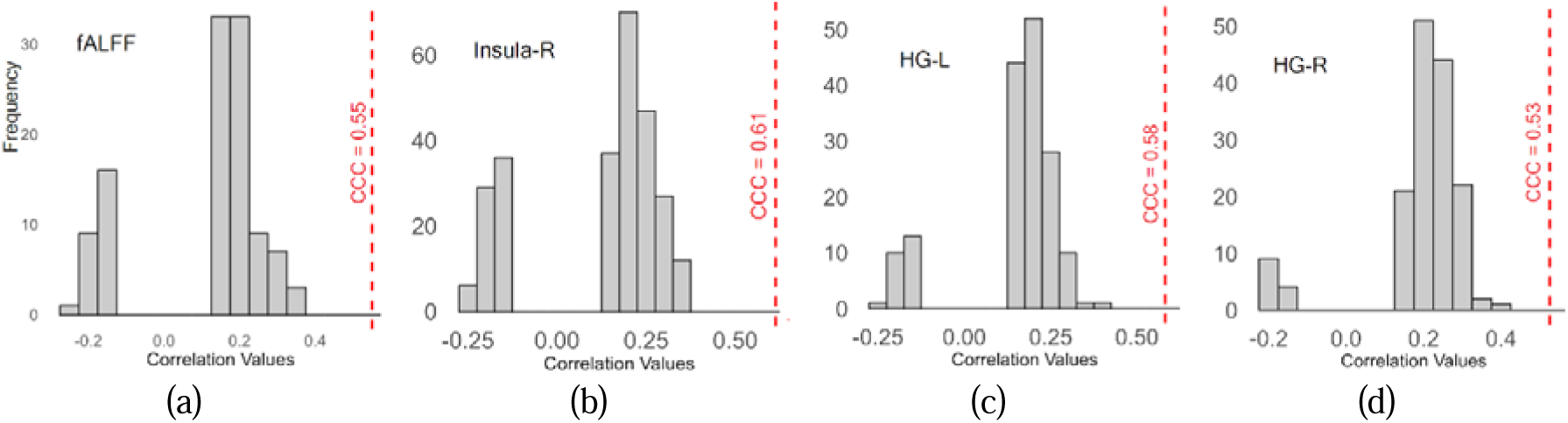
Comparing correlation of individual structural and functional metrics with the correlation of four corresponding admissible combined structural and combined functional (CS-CF) features. The bar plots illustrate the histogram of linear correlation coefficients of all individual structural and (a) fALFF (b) FC (seed: insula-R), (c) FC (seed: HG-L), (d) FC (seed: HG-R) metrics. The vertical red dotted line in each diagram specifies the canonical correlation coefficient (CCC) of the corresponding admissible CS-CF features. HG: Heschel’s gyrus, L: left, R: right

The details of the significant correlations of individual structural and functional metrics of all PSRs (excluding operG, AG, MTG in both hemispheres) are presented in Fig. S2 to Fig. S12 in Supplementary Materials. The heatmaps in Fig. S2(a) to Fig. S12(a) show the significant S-F correlation for individual structural metrics. Figures S2(b) to S12(b) illustrate the histogram of correlations between individual metrics in comparison to CCC for combined features. Additionally, Fig. S2(c) to Fig. S12(c) present the regression line for the proposed CS and CF features. According to the correlations between individual metrics, myelination index is the most correlated structural metric to CF features.

The correlation coefficients between individual structural and functional metrics in Fig. S2 to Fig. S12 are mostly consistent with previous studies [16]. The previous studies demonstrated the positive correlation between myelination and FC, which shows the development of myelination causes stronger FCs [11]. Various factors including neuronal migration to gray matter and programmed cell death with different timings for each region cause various postnatal regional developmental patterns of surface morphology [61]. Consequently, different patterns of S-F relation are expected when considering all regions. Negative correlations between cortical thickness and FC are mostly consistent with previous findings [11]. The respective positive and negative relation between FC and sulcal depth and curvature are in line with previous studies on cortical folding. For example, the correlation between sulcal depth and long-range FC is positive, while its relation with local FC is negative [62].

### 3.2. Canonical weights configuration in admissible CS-CF features

The brain maps of canonical weights corresponding to fALFF and myelination of 16 PSRs in the admissible CS-CF features from the first experiment are shown in Fig. 3(a). The corresponding canonical correlation coefficient is 0.55. Considering the canonical weights in Fig. 3(a) and the details presented in Table S4, the fALFF of insula and HG in both hemispheres contribute mostly in CF feature. The myelination of insula in both hemispheres, MTG-R, HG-L, trianG-R, AG-R and STG-R mostly contributed in CS feature.

**Fig. 3.**
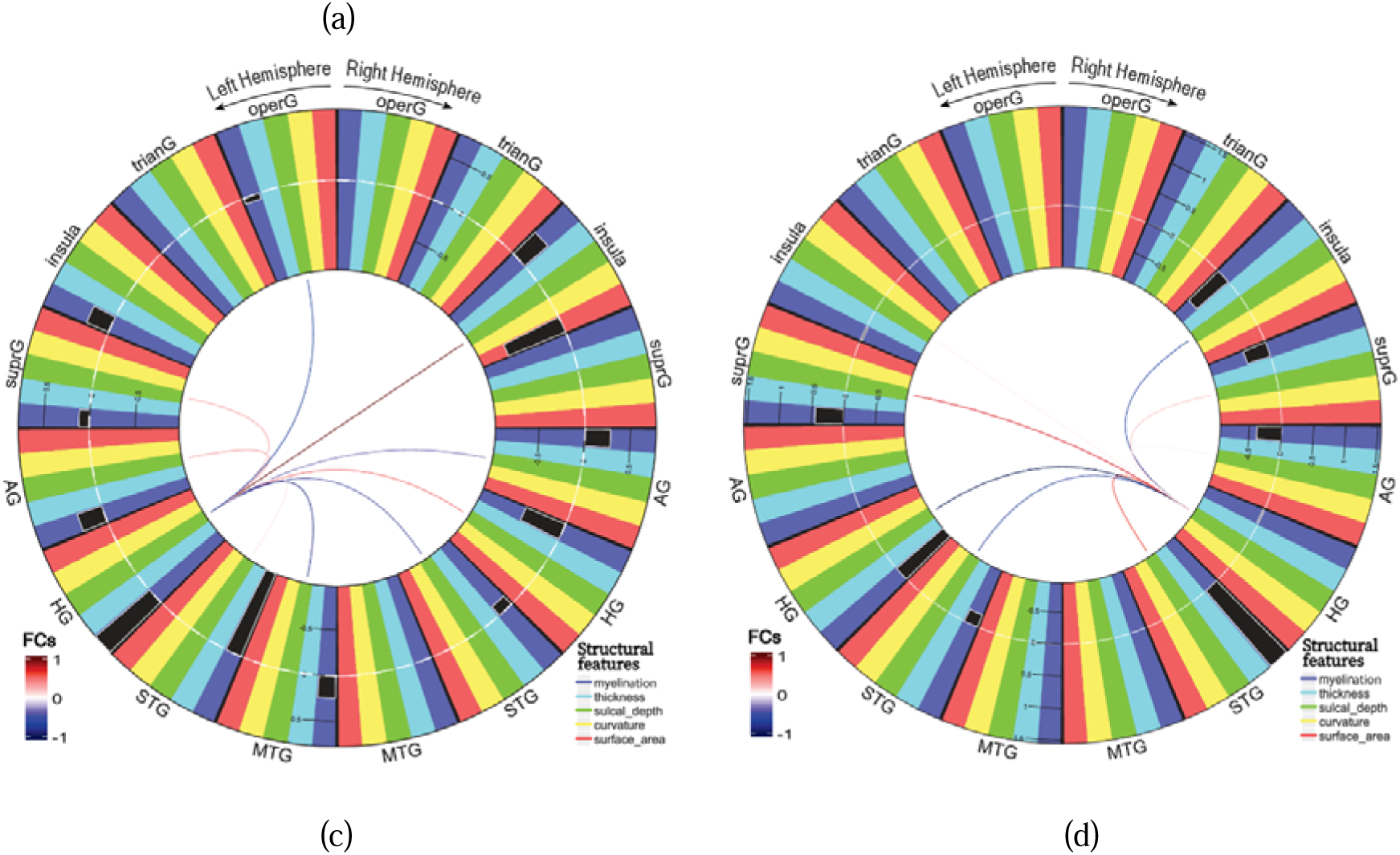
Canonical weights of each metric from 16 PSRs in CS-CF features. (a) Brain maps illustrate the canonical weights obtained by the first experiment using fALFF as functional features. As illustrated, the myelination is the only structural metric contributed in CS feature. (b-c) Canonical weights obtained by the second experiment using FC between the designated PSR and remaining PSRs. The arcs connecting (b) insula-R, (c) HG-L and (d) HG-R to the remaining PSRs, represent their FCs. The connection color specifies the canonical weight of the FC which takes part in CF feature. The black bars inside the radial color bars around each connectogram shows the canonical weights of structural features. trianG: triangular gyrus, operG: opercular gyrus, suprG: supramarginal gyrus, AG: angular gyrus, STG, MTG: medial temporal gyrus, HG: Heschel’s gyrus, L: left, R: right

Furthermore, the canonical weights for FCs originating from the insula-R (CCC=0.61), HG-R (CCC=0.58) and HG-L (CCC=0.53), in the second experiment are depicted using connectograms in Fig. 3(b-d). In these connectograms, 16 PSRs are represented as nodes. Each node is represented by five colored segments, each corresponding to a different structural feature. The black bars inside the radial color bars around each connectogram show the canonical weights of correlated structural features. The PSRs in the right and left hemispheres are allocated respectively to the right and left semicircles in this diagram. The arcs connecting pairs of nodes in the inner circle represent FCs between PSRs, and their colors specify the corresponding FC value. The FC between inula-R and the following regions contribute in the corresponding CF feature in order of their contributions: operG-L, insula-L, AG-L, suprG-L and trianG in both hemispheres. The canonical weights show the myelination contribution of inula-L, trinaG-L, supr-R and operG in both hemispheres are high in the corresponding CS feature (Fig. 3(b) and Table S5). The FC between HG-L and insula-R, MTG-L and opercG-L, as well as myelination of STG-L and HG in both hemispheres, and surface area of inula-R contribute mostly in the corresponding CS-CF features (Fig. 3(c) and Table S6). As shown in Fig. 3(d) and detailed in Table S6, the FC between HG-R and HG, superG and STG in left hemisphere contribute mostly in CF feature. The myelination of HG and superG in left hemisphere and STG in right hemisphere have higher canonical weights in the corresponding CS feature.

We further investigated the redundancy of the structural and functional metrics which have been discarded by the extracted CS and CF features. For this purpose, we computed the Pearson correlation coefficient between individual structural metrics and CF feature, as well as between individual functional metrics and CS feature. The correlation coefficients are shown in Fig. S13 and Table S8 to S11 in Supplementary Materials. As expected and illustrated in Fig. 3 and Fig. S13, certain morphological metrics did not contribute to CS feature and they are less correlated with CF features in contrast to myelination which contributes in CS feature. In the first experiment, for example, the myelination of HG in both hemispheres, AG-L and MTG-L are highly correlated individual structural metrics with CF features. While MTG-L is a discarded region, its structural and functional metrics have significant correlation with CF and CS features, respectively. Given the limited number of significant correlation coefficients between structural metrics and CF features, the following attributes were excluded despite demonstrating significant correlations: thickness of AG-L, sulcal depth of MTG-L, as well as the curvature and surface area of MTG-L and HG-L. There are additional examples in the second experiment, like when HG-L was used as a seed; surface area of STG-R and AG in both hemispheres, and curvature of AG-R are other discarded metrics with significant correlation with CF feature, while the myelination of these regions has contributed in CS and have correlation with CF feature. In this analysis, other discarded PSRs with considerable correlation between their myelination and CF features are opercG-R and trianG in both hemispheres. The surface area of these regions have significant correlation with CF features, as well.

## 4. Discussion

The value of the S-F approach to study brain aging and disorders have been demonstrated by recent investigations. Zimmermann et al. [63] illustrated that the age-related S-F association can provide additional insights into brain aging beyond what can be revealed by each modality alone. Furthermore, Rudie et al. [64] employed a similar approach to show that social and communication symptom severity of autism spectrum disorder (ASD) are associated with an imbalance of global and local efficiency between structural and functional networks. Accordingly, the S-F approaches are potentially promising to provide predictive features for early diagnosis or prediction of brain and behavioral disorders. Jiang et al. [65] investigated the decrease in S-F coupling to predict suicide attempts in subjects with bipolar disorder. Likewise, Wen et al. [66] discovered the link between global increase in S-F coupling as well as within specific brain modules and decreased dynamic brain function in Tourette Syndrome. They also found that the S-F correlation could serve as a biomarker for early diagnosis of Tourette Syndrome.

Establishment of the S-F atlas of normal brain development may help prediction or early diagnosis of brain or behavioral disorders. The early postnatal period is crucial for investigating brain development, since the brain undergoes fundamental changes and any deviation from normal growth may yield to subsequent structural or functional disorders. A few S-F studies tried to model the neonatal brain development in preterm and term subjects using individual features [11,12,16,17,33,34]. The objective of this study was to improve the present understanding of the S-F relationships by examining CS-CF features as an alternative to individual S-F features using a large sample size of term neonates. Similar to most of the previous early neurodevelopmental studies [33,34], this study was focused on PSRs because they can be investigated via passive responses of the subjects to auditory and language stimulations.

A limited number of S-F studies utilizing neonatal rs-fMRI based on diverse individual structural and functional metrics have been conducted. These investigations identified whole-brain S-F correlations prior to regressing out the age [11,15,16]. Subsequent to controlling for age, the S-F relationship was observed primarily between a restricted set of features [16,17], with particular challenges noted for term neonates compared to preterms [17]. Daneshvarfard et al. [33] found no significant S-F relationships after regressing out the age in auditory regions. Another study indicated S-F correlation only in one neonatal PSR after controlling for the age [36]. This may be attributed to using individual structural and functional features, as suggested in a way by the comparison between direct and network-based microstructural connectivity and FC relationships in S-F analysis [17]. With this perspective, we examined the S-F relationship using combined structural and combined functional features after regressing out the age. The correlation observed in this study from the combined structural and functional features was not only significant but also surpassed the correlation obtained between function and structure in the previous whole brain studies [11,16,17].

The association between myelination as well as morphological features with different brain functions is investigated in early stages of life. Compared to morphological features, the microstructural metric showed the strongest relationship to the spatial variations in cortical gene expression throughout gestation [67]. Larivière et al. [11] showed T1w/T2w ratio is significantly correlated with principal functional gradient in 40 term neonates, while cortical thickness is weakly correlated with the second principal functional gradient of the posterior to anterior regions. Similarly, a regression analysis demonstrated the relation between the neonatal function of the brain and myelination is stronger than morphological features [16]. In our study, the correlations between individual features confirmed the strong coupling between myelination and functional features. Furthermore, the canonical components in our age-controlled S-F analysis in PSRs, showed stable and significant relationships mostly between myelination metrics and functional ones including fALFF or FCs.

The postnatal dramatic growth of neurons reflects itself in changes of the morphological metrics like surface area and cortical thickness [5,68] which is correlated with the function of the brain. Our study demonstrated limited contribution of morphological metrics such as surface area in CS features which is in line with previous results and neurobiological evidences [11,16]. We found also weak correlations between individual morphological metrics and CF features which confirm previous findings. In contrast, our results show the effect of myelin sheath around axons on the brain’s function is prominent to the effect of morphological features and myelination index contains some shared information regarding the relationship between morphological and functional features. However, the discrepancy of the correlation between function and cortical thickness in our results and the results presented in [11], may be due to regressing out the age.

Our findings on the relation between fALFF and myelination is in accordance with [16], while the significant correlation obtained between FC and CS features disapprove their results. From a different view point, Gondová et al. [17] showed that network based information of FC and microstructural connectivity establish the S-F relation better than their direct utilization. This may confirm the potential of our CS features in establishing relation with combined FCs.

To be more specific about the contribution of PSRs in combined features, the FCs of inula-R, HG in both hemispheres with other PSRs, relate to the myelination index. The insula is a major afferent hub that receives extensive viscero-autonomic feedbacks [69,70]. Its role as a hub region is supposed to begin during neonatal period [71]. This area belongs to the salient network that establishes the coupling with the auditory network in preterm neonates [72]. Accordingly, our results show the insula as a seed has more stable FCs with other PSRs than other selected seeds that is HG in both hemispheres. Additionally, the canonical weights show insula and HG have contributions in all CS-CF features, including the admissible combined as well as the other combined features. There is a bilateral FC between HG and auditory, sensorimotor, cingulo-opercular, and visual networks in old adults during rest [73]. The FCs between HG and cingulo-opercular network support auditory attention processes. They are correlated with hearing ability and uncorrelated with age or cognitive performance [73]. While our CS-CF analysis is limited to the PSRs, the contribution of HG in both CS and CF and specially the connection between HG-L and operG-L in CF feature are shown. More analysis between HG and other brain regions are needed to investigate the S-F coupling of the regions which support auditory attention during early stages of postnatal development. The direct investigation of regional FC and microstructural connectivity [17] showed no significant coupling in inter-hemispheric homotopic and non-homotopic connections in term neonates. Nevertheless, network-based analysis in [17] demonstrated a significant S-F coupling which suggests more complex S-F interactions in term neonates which require further investigation to fully understand their underlying mechanisms. The visual inspection of connectograms in Fig. 3 suggests the prominent incorporation of inter-hemispheric FCs in CF feature in relation with CS feature during postnatal development in term neonates. The agreement between our findings and those of network-based analysis in [17] may stem from our combined features approach.

Using the sparse CCA method to select features has two advantages. First, it enables us to explore multi-dimensional S-F space to find correlations stronger than what individual S-F features provide in perisylvian region [33,34]. Second, it looks for a sparse and optimal subspace in which a combination of selected features establishes the S-F relationship. In our approach, like [60], the significant components and stable weights were selected based on CCs and bootstrapping tests respectively. However, the features removed by bootstrapping tests may reduce the correlation between the remaining combined features. As removing some weights from the selected components required further analysis, consideration of canonical weights significancy [74] while computing canonical components, can help address this issue in future studies.

The relatively small sample size of neonates compared to the dimension of structural and functional metrics was one of the limitations of our study. Despite the use of sparse parameters to mitigate this limitation, a larger number of participants would improve the generalization of the resulting features. Additionally, the inclusion of diverse populations across various ages allows for a more comprehensive investigation of S-F relations.

Our study being focused on examining the S-F relationship in normal subjects, we enrolled only term infants. It is crucial to consider the infants’ health beyond the immediate post-birth period and conduct longitudinal studies. The availability of longitudinal health data for the neonates participating in our study would prevent infiltration of suspicious subjects and improve our model of the S-F relationships in a healthy brain. Obviously, the excluded subjects may serve as test samples in order to assess the prognosis ability of the created S-F feature.

Like the previous neonatal S-F studies [12,15,33,34], we did not take into account the gender of neonates. The sex difference of the neonatal brain is believed to be limited, because of the restricted differential sex chromosome expression and weak production of sex hormones during the early stage of development. Nevertheless, inconsistent results of structural and functional studies show limited sex-based differences in neonatal brain [5,75]. There is no evidence on the neonatal sex differences of cortical thickness and global gyrifications [76–79]. Sex-related differences of microstructural measures are still unknown [5,80]. Some regional sexual dimorphism in brain size have been reported in the early stages of development [81–83], while there are no significant sex-based differences of the global brain volume [84,85]. Recently, some perinatal sex-related differences of FC patterns were demonstrated [86–88]. Further investigations may help to reveal the effect of gender on neonatal CS-CF features.

This study developed CS-CF features, considering the linear S-F association. However, recent studies demonstrated the potential of non-linear models like deep neural networks to reveal more complex S-F relationships [89,90].

Consistent with former studies [11,16], our findings indicate that the myelination index as a microstructural metric shows higher correlation with FC than morphological metrics. Other microstructural metrics (e.g., fractional anisotropy, mean diffusivity) based on diffusion MRI may be considered for correlation with FC [91,92]. Likewise, the relation between FC and structural connectivity can be considered in future works. Moreover, investigating the relation between structural metrics and other local functional ones like regional homogeneity (Reho) would be advantageous. Overall, future studies using non-linear models and a variety of structural and functional metrics and modalities may help to achieve a comprehensive CS and CF features during brain development.

Some structural and functional asymmetries in perisylvian sub-regions have been reported by previous studies during the perinatal period [93–97]. It seems that there are some differences in S-F relationships between the left and right perisylvian sub-regions in our study. Further studies should be conducted to investigate the significance of laterality of the neonatal S-F associations.

In this study, we used concurrent changes of structure and function of subjects to extract the S-F couplings. Further studies may be performed to investigate correlations between asynchronous structural and functional features of different ages, in order to explore the lead/lag development of the structure versus function.

## 5. Conclusion

In this study, we proposed CS-CF features for the perisylvian sub-regions in neonates by integrating various metrics extracted from functional and structural MRI data using CCA. This method addresses the limitation of having a small number of neonates relative to the number of features by leveraging a sparse combination of features. Statistical analysis revealed the contribution of myelination and FCs, consistent with the previous studies. Moreover, the relationships between myelination and previously reported FC of some perisylvian sub-regions were demonstrated using CS-CF features. Further studies may explore nonlinear or asynchronous relationships between structure and function. Future work should also investigate the application of CS-CF features in the diagnosis and prognosis of neonatal brain disorders.

## Acknowledgement

This work was supported by Cognitive Sciences and Technologies Council (COGC), Iran [grant numbers 96P97, 9747].

